# A Reproducible Supermicrosurgical Model of Cervical Lymphatic-Venous Anastomosis in a Transgenic Alzheimer’s Mouse: Technical Translation, Proficiency, and a Validated Learning Curve

**DOI:** 10.1101/2025.08.11.657918

**Authors:** Renpeng Fang, Lei Jin, Jianlong Wu, Qingping Xie, Ming Xiao, Xiaodong Yang, Hongrui Lu, Maximilian Kueckelhaus

## Abstract

**Background:** Cervical Lymphatic-Venous Anastomosis(LVA) holds therapeutic potential for Alzheimer disease (AD). However, a validated AD animal model for preclinical investigation remains lacking. This study aimed to establish and standardize LNVA, as a form of lymphatic-venous Anastomosis, in a murine AD model; to quantitatively assess its surgical learning curve; and to create a platform for mechanistic exploration and translational research.

**Methods:** We established and standardized a deep cervical lymph node-to-vein anastomosis (DC-LNVA), a specific type of lymphatic-venous anastomosis (LVA), in rodents. A two-stage training program was applied to a junior surgeon: 60 DC-LNVA procedures were performed in 30 small Sprague-Dawley rats (50–60g), followed by 20 bilateral anastomoses in 10 male 5XFAD mice. Surgical proficiency was evaluated using procedure time, UWOMSA scoring, and CUSUM analysis. Anatomical parameters were assessed with ImageJ. Anastomotic patency was confirmed via ICG lymphangiography. Intranasal Evans Blue was used to trace cranial distribution pathways.

**Results:** SD rats had significantly larger external jugular veins (0.376 ± 0.013 mm vs. 0.228 ± 0.011 mm) and DCLNs (1.359 ± 0.084 mm^2^ vs. 0.333 ± 0.022 mm^2^) than mice (P < 0.0001). Proficiency was achieved after 27 anastomoses in rats and 10 in mice. ICG confirmed 100% patency in 9 surviving 5XFAD mice. Evans Blue suggested a nasal–cranial pathway for tracer distribution.

**Conclusion:** We developed and validated a cervical lymphatic-venous bypass (LVA) in transgenic AD mice. This platform complements existing loss-of-function approaches and offers a foundation for mechanistic and translational research into brain lymphatic clearance.

## Introduction

The discovery of the glymphatic system and meningeal lymphatic vessels have fundamentally shifted our understanding of how the brain clears metabolic waste. ^1–3^ This has opened new avenues for investigating Alzheimer’s disease(AD), where impaired waste clearance may contribute to disease progression.^4,5^ Research on the brain’s lymphatic system has rapidly gained momentum in recent years, emerging as a key frontier in Alzheimer’s disease research.^6^ In preclinical studies, surgical ligation of the deep cervical lymph nodes (DCLNs) is commonly employed as a loss-of-function model to investigate downstream pathological changes in brain lymphatic drainage.^3,7^ For example, Xiao et al. reported that DCLNs ligation accelerates amyloid-beta accumulation and exacerbates Alzheimer’s disease(AD) pathology.^8^ More recently, Kim et al. demonstrated that this surgical ligation of DCLNs affects IL-6 dependent signaling, leading to an imbalance in cortical excitatory and inhibitory and resulting in memory deficits.^9^

While such loss-of-function models have provided valuable insights into the role of DCLNs, a complementary gain-of-function approach is needed to enable bidirectional validation and enhance methodological rigor. Several pharmacologic strategies, such as the use of prostaglandin F2α and VEGF-C, have been explored to promote cervical lymphatic drainage.^9,10^ However, their systemic effects and lack of anatomical specificity limit their ability to precisely modulate DCLNs function. To date, no surgical gain-of-function model has been established to supplement existing loss-of-function studies.

In parallel with these preclinical considerations, the therapeutic potential of a surgical “gain-of-function” strategy has recently been translated into clinical practice. Our group reported the initial application of deep cervical lymph node-venous anastomosis (LNVA)—a mature microsurgical technique—to patients with Alzheimer’s disease, with the aim of enhancing cerebral lymphatic outflow.^11–13^ This novel application has since generated considerable interest within the international lymphatic and neurosurgical communities, leading to its growing adoption by numerous medical centers, promising preliminary outcomes reported by several centers, and the initiation of international clinical trials.^14–16^

This rapid clinical translation, along with the encouraging early results, amplifies the urgent need for a well-characterized and reproducible preclinical platform to investigate the underlying therapeutic mechanisms. We propose that the true utility of such a surgical model lies in its accessibility—its feasibility must extend beyond elite experts to proficient junior surgeons to facilitate widespread adoption in research. While our group previously reported a foundational model in rats,^17^ advancing this technique to the mechanistically critical transgenic mouse models has been hindered by a formidable supermicrosurgical barrier. Therefore, the present study was designed to systematically overcome this challenge through an innovative, staged training protocol, ultimately establishing and validating the technique in a transgenic AD mouse model to provide a robust and reproducible platform for future research.

## Methods

### Ethics Statement

All procedures were approved by the Institutional Animal Care and Use Committee (IACUC).All procedures adhered to national guidelines for the care and use of laboratory animals. Rats (training model) Male Sprague-Dawley (SD) rats, 3—4 weeks old, 50—60 g, supplied by Hangzhou Qizhen Laboratory Technique Co. (Hangzhou, China).

Mice (application model) Male 5×FAD transgenic mice,^18^ 24—32 weeks old, 20—35 g, obtained from the Laboratory of Neurodegeneration, Nanjing Medical University (Nanjing, China). Animals were housed in a controlled environment (22 ± 2 °C, 50 ± 10 % humidity, 12-h light/dark cycle) with ad libitum access to food and water. All animals were acclimatized for ≥ 7 days before any intervention. The work has been reported in accordance with the ARRIVE guidelines (Animals in Research: Reporting In Vivo Experiments).^19^

### Study Design

The project comprised two sequential phases led by a single junior microsurgeon (three years’ experience) under the supervision of two senior microsurgeons: Training Phase (SD rats, n = 30): Master cervical lymph node-to-venous anastomosis (LNVA) on a transitional model whose vessel calibers approximate those of mice. Learning-curve analysis and definition of a standard operating procedure (SOP). Application Phase (5×FAD mice, n = 10): Apply the rat-refined SOP to establish a gain-of-function LNVA cohort for downstream Alzheimer’s-disease research.

### Surgical Training and Skill Assessment

Video-recorded anastomoses were independently and blindly scored by the two senior microsurgeons using the University of Western Ontario Microsurgical Skills Acquisition (UWOMSA) instrument (scale: 1 = novice to 5 = expert, 0.1-point resolution).^20^ The mean of both ratings constituted the final score for each procedure. Anastomotic patency was verified intraoperatively by direct visualization and, when indicated, by indocyanine green (ICG) lymphangiography.^21^ Instances of non-patency triggered immediate expert feedback, including analysis of technical errors and corrective guidance.

### Surgical Procedure

Both Sprague-Dawley (SD) rats and 5xFAD transgenic mice were anesthetized via intraperitoneal (IP) injection of a ketamine (100 mg/kg) and xylazine (10 mg/kg) solution. Ophthalmic ointment was applied to prevent corneal desiccation. Core body temperature was maintained at 37.0 ± 0.5°C using a thermostatically controlled heated surgical platform.

The surgical technique was adapted from established human clinical procedures and modified for rodent anatomy.^22,23^ A single midline cervical incision was employed to provide bilateral access to the DCLNs,**(Figure 1A) s**implifying the procedure while enabling bilateral anastomoses. Animals were secured supine after anesthesia. The surgical protocol was consistent for both rats and mice due to comparable cervical anatomical landmarks. The platysma muscle was incised longitudinally along the midline to expose deeper cervical structures. The sternocleidomastoid (SCM) muscle was identified and retracted medially to reveal the surgical field. Deep to the SCM, a distinct cluster of fatty tissue encapsulating the DCLNs was visualized. **(Figure 1B)** The targeted DCLN was meticulously isolated using supermicrosurgical techniques, emphasizing preservation of surrounding afferent and efferent lymphatic vessels and nodal integrity. Complete skeletonization of the DCLN was avoided, following the technique detailed by Chen et al.^23^ Instead, the node was mobilized with a small margin of its associated adipose and connective tissue, akin to creating a ‘lymphatic-fatty flap’ or ‘lymphatic pedicle’. This aimed to protect delicate lymphatic connections to the node and maintain optimal viability of the nodal tissue selected for anastomosis. The EJV ipsilateral to the selected DCLN was carefully isolated from its fascial sheath over a suitable length. **(Figure 1C)** Notably, in mice, the EJV generally possesses a larger caliber than the internal jugular vein in this anatomical region, making it the preferred target vein. Temporary occlusion of the EJV segment was achieved using microvascular clamps (proximal and distal to the planned venotomy site). In mice, where the operating space was more confined, an alternative occlusion method involving a temporary slipknot ligature with an 8-0 silk suture was occasionally employed to ensure precise blood flow control. The isolated DCLN was prepared for anastomosis by carefully bisecting it transversely with micro-scissors at approximately its midpoint. This maneuver exposed the internal lymphatic architecture of the node. **(Figure 1D)** The bisection was performed aiming to preserve afferent lymphatic inflow to the proximal nodal segment that was selected for anastomosis, thereby creating a viable, patent surface rich in exposed sinuses.

**Figure 1.**
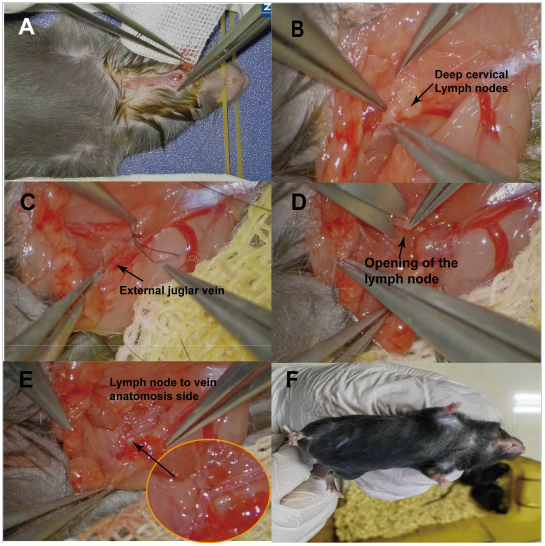
Surgical procedure (A) A midline cervical incision (1.0–2.0 cm) is made, and the platysma is split. (B) The sternocleidomastoid muscle is retracted to expose deep cervical lymph nodes (DCLNs) within surrounding fat. (C) The external jugular vein (EJV) is isolated and temporarily occluded using a microvascular clamp (rat) or 8-0 silk slipknot (mouse). (D) The DCLN is bisected while preserving afferent lymphatics; a venotomy is created to match the cut surface. (E) Anastomosis is performed using 10-0 (rat) or 11-0/12-0 (mouse) nylon sutures. The contralateral side is treated similarly, and incisions are closed in layers. (F) One month postoperatively, mice showed good survival and spontaneous suture loss, indicating complete wound healing.

Subsequently, a longitudinal venotomy, sized to adequately match the transected surface diameter of the lymph node segment, was created in the anterior wall of the prepared EJV using microscissors.An end-to-side (lymph node-to-vein) anastomosis or end-to-end (lymph node-to-vein branch) anastomosis was performed between the prepared transected surface of the DCLN segment and the venotomy in the EJV. **(Figure 1E)** For procedures in rats, 10-0 nylon sutures were predominantly used. For the more delicate anastomoses in mice, 11-0 or, occasionally, 12-0 nylon sutures were employed. Upon completion of the anastomosis, the microvascular clamps or temporary ligature were released. Initial patency was grossly assessed by observing for satisfactory blood refilling of the venous segment and the absence of significant leakage from the anastomosis site, along with the appearance of the anastomosed lymph node. The contralateral surgery was then performed using the same meticulous technique. Finally, the platysma muscle and skin incisions were closed in layers using 5-0 absorbable sutures.

### Indocyanine Green (ICG) Lymphangiography

To assess lymphatic-venous connectivity, 0.05 mL of a 0.25 mg/mL Indocyanine Green (ICG; Ruidu®, Dandong Yichuang Medical Company, China; from a 25 mg vial) solution was administered via unilateral nasal instillation, targeting the nasal mucosa. **(Figure 2A**,**B)** The injection was performed as superficially as possible, stopping upon sensing slight resistance to avoid deeper penetration and unintended vascular entry. A 30-minute waiting period was observed to allow for ICG distribution within the lymphatic system before imaging. ICG was conducted using the integrated fluorescence system of the surgical microscope. **(Figure 2C**,**D)**

**Figure 2.**
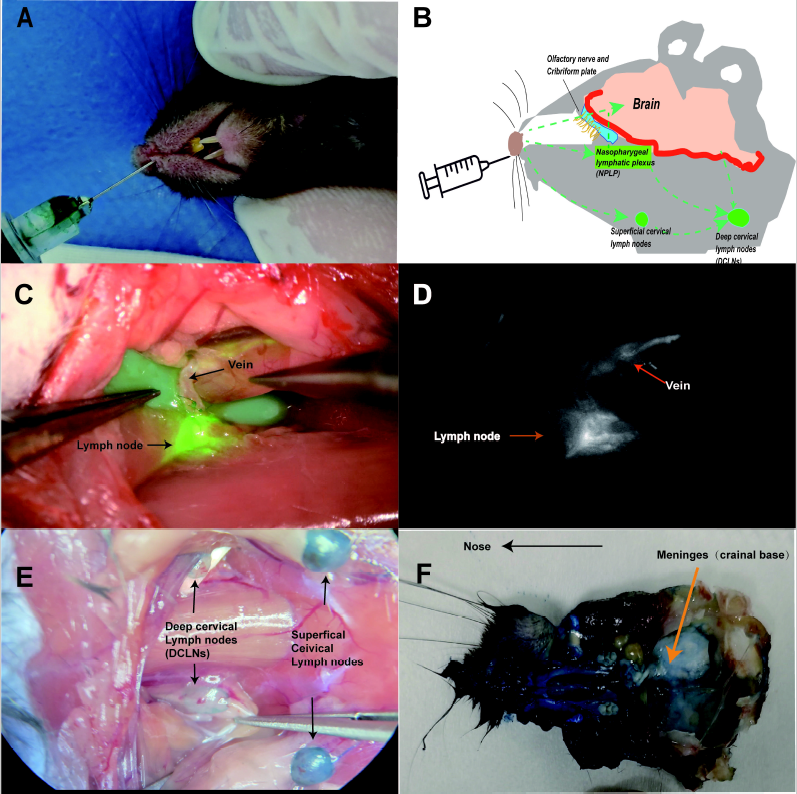
ICG injection (A) Schematic of intranasal tracer injection in mice. (B) Hypothetical diagram illustrating nasal lymphatic drainage pathways following mucosal administration. (C) Intraoperative image of indocyanine green (ICG) fluorescence overlaid with surgical field under microscopy. (D) Real-time ICG lymphangiography demonstrating tracer flow from the injection site. (E) Following Evans Blue injection, superficial cervical lymph nodes show early uptake, followed by bilateral deep cervical lymph nodes. (F) Postmortem dissection reveals Evans Blue staining in the nasal cavity, nasopharyngeal lymphatic plexus, and dura mater adjacent to the skull base.

### Surgical Equipment, Instruments, and Sutures

All microsurgical procedures on Sprague-Dawley (SD) rats were performed under a Zumax OMS3500 surgical microscope (Zumax Medical, Suzhou, China). For the more delicate procedures on 5xFAD transgenic mice, a Zeiss KINEVO 900 robotic visualization system (Carl Zeiss Meditec, Oberkochen, Germany) was utilized, providing enhanced magnification and illumination. Supermicrosurgical instruments, including forceps with tip diameters of 0.10 mm and 0.15 mm, were obtained from Micro Fox (Shanghai Guli Science and Technology Co., Ltd., Shanghai, China). Anastomoses in rats were performed using 10-0 and 11-0 nylon sutures (Ningbo Lingqiao Medical Apparatus Co., Ltd., Ningbo, China). For mice, procedures utilized primarily 12-0 nylon sutures (CrownJun, Japan), with 11-0 nylon sutures (Ningbo Lingqiao Medical Apparatus Co., Ltd., Ningbo, China) also being used occasionally.

### Data Collection and Processing

Comprehensive data were collected during all surgical procedures for key step documentation, morphometric analysis, and learning curve construction.

Intraoperative images and videos were systematically captured for documenting surgical steps, morphometric analysis, and video-based UWOMSA skill assessment. External jugular vein (EJV) diameter and deep cervical lymph node (DCLN) planar area were measured from calibrated intraoperative images of SD rats and 5XFAD transgenic mice using ImageJ software. Microsurgical forceps tip diameters (0.10 mm and 0.15 mm) served as calibrated references. For each of 60 anastomoses in SD rats and 20 in 5XFAD mice, surgical time (left and right sides separately) and UWOMSA scores (0–5 scale) were meticulously recorded sequentially to generate datasets for learning curve analysis.

### Statistical Analysis

Statistical analyses were performed using R software (Version 4.4.2; R Core Team, Vienna, Austria). Data are graphically presented as mean ± standard error of the mean (SEM). A p-value < 0.05 was considered statistically significant. In figures, the notation “****” indicates a p-value < 0.0001. Differences in anatomical parameters (External Jugular Vein diameter and Deep Cervical Lymph Node area) between SD rats and 5XFAD transgenic mice were analyzed using two-tailed unpaired Student’s t-tests. Learning curves for surgical time and UWOMSA scores were plotted against the cumulative number of anastomoses, with trendlines generated using smoothing algorithm added to visualize skill progression. To objectively determine proficiency, Cumulative Sum (CUSUM) analysis was applied to both surgical time and UWOMSA scores. CUSUM values were calculated for each sequential procedure relative to the overall mean for that parameter. The resulting CUSUM data were then fitted with polynomial regression models (cubic regression for SD rat data, n=60 anastomoses; quadratic regression for 5XFAD mouse data, n=20 anastomoses). The vertex of the fitted CUSUM curve was identified as the point at which proficiency was achieved.

## Results

To establish a baseline for the surgery procedure, key anatomical parameters were measured in both Sprague-Dawley (SD) rats and 5XFAD transgenic mice. The External jugular vein (EJV) diameter was significantly larger in SD rats (0.376 ± 0.013 mm, n=60 measurements) compared to 5XFAD mice (0.228 ± 0.011 mm, n=20 measurements; P < 0.0001, **Figure 3A**). Similarly, the planar area of the deep cervical lymph nodes (DCLNs) was substantially greater in SD rats (1.359 ± 0.084 mm^2^, n=60 measurements) than in 5XFAD mice (0.333 ± 0.022 mm^2^, n=20 measurements; P < 0.0001, **Figure 3A**). These dimensional differences highlight the increased technical demand anticipated for procedures in the smaller mouse model.

**Figure 3.**
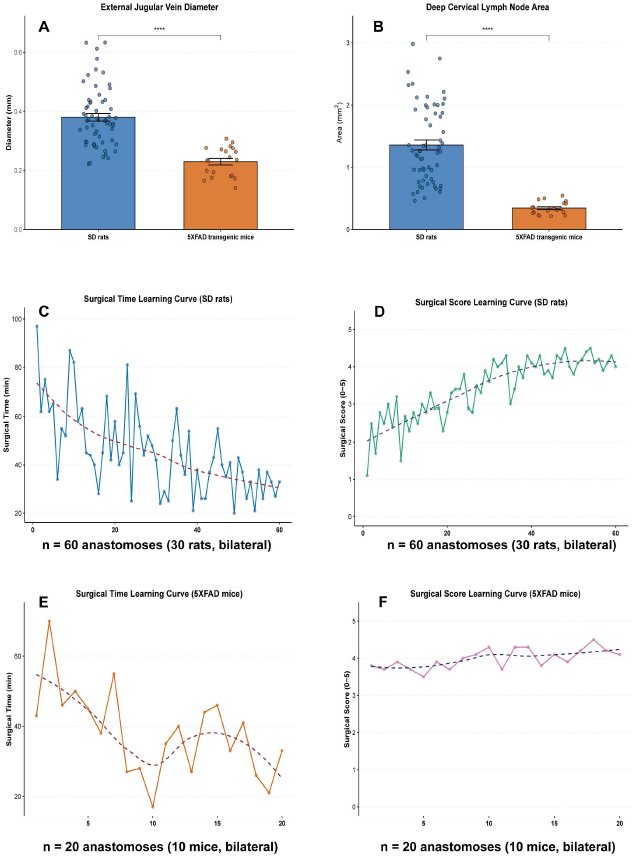
Anatomy and Surgical Learning Curves (A) External jugular vein diameter was significantly larger in SD rats compared to 5×FAD transgenic mice (****P < 0.0001). (B) Deep cervical lymph node area was also significantly greater in SD rats (****P < 0.0001). (C–D) Learning curve in SD rats (n = 30 rats, 60 anastomoses): (C) Surgical time per anastomosis decreased over time; (D) surgical proficiency scores progressively improved based on UWOMSA evaluation. (E–F) Learning curve in 5×FAD mice (n = 10 mice, 20 anastomoses): (E) Operative time showed a downward trend; (F) surgical scores remained stable with minor fluctuations, reflecting adaptation from prior rat training. Dotted lines indicate polynomial trendlines.

A junior surgeon, with prior general microsurgical experience, underwent a structured training program performing 60 surgery procedures in 30 SD rats (bilateral anastomoses). Analysis of the learning curve revealed a progressive improvement in surgical efficiency and skill. Surgical time demonstrated a significant decrease with cumulative experience, from an initial average exceeding 80 minutes to consistently below 40 minutes in later procedures **(Figure 3C)**. Concurrently, surgical skill scores, assessed via the UWOMSA scale, steadily increased, indicating enhanced procedural quality and proficiency **(Figure 3D)**.Cumulative Sum (CUSUM) analysis was employed to objectively determine the inflection point signifying attained proficiency. For surgical duration in SD rats, the CUSUM curve fitted with a cubic regression model (R^2^ = 0.9638) identified the proficiency vertex after approximately 27 anastomoses **(Figure 4A)**. For surgical skill scores, a cubic regression fit (R^2^ = 0.9916) of the CUSUM data indicated that proficiency was achieved at approximately 26 anastomoses **(Figure4B)**. These analyses, supported by high coefficients of determination, suggest that consistent proficiency for the surgery in SD rats was reached after approximately 26-27 anastomotic procedures.

**Figure 4.**
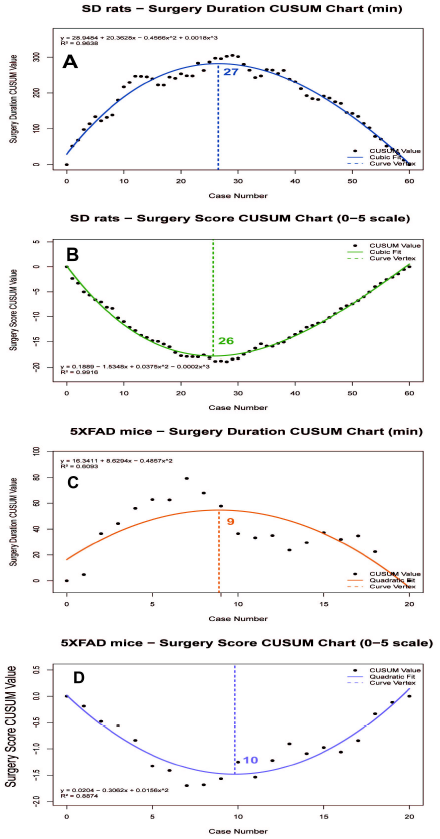
CUSUM analysis (A) Cumulative sum (CUSUM) analysis of surgical duration for SD rats (n = 60 anastomoses) shows a peak at case 27, indicating the learning curve vertex. (B) CUSUM chart of surgical performance scores for SD rats identifies a minimum at case 26, consistent with plateaued skill acquisition. (C) In 5×FAD mice (n = 20 anastomoses), the CUSUM chart of surgical duration peaks at case 9, marking procedural stabilization. (D) CUSUM analysis of surgical scores in 5×FAD mice shows a minimum at case 10, suggesting early adaptation post rat-model training. Solid lines represent fitted curves (cubic or quadratic), and vertical dashed lines indicate curve vertices.

Following proficiency attainment in SD rats, the surgery technique was transferred to 5XFAD transgenic mice (10 mice, 20 bilateral anastomoses). A marked transfer of skill was evident, with initial surgical times in mice being considerably shorter and baseline skill scores higher than those observed at the commencement of rat training. The learning curve for surgical time in 5XFAD mice also showed a decreasing trend with practice **(Figure 3E)**, and surgical skill scores further improved, rapidly reaching a high level of competence **(Figure 3F)**. CUSUM analysis confirmed this accelerated learning in the mouse model. The proficiency vertex for surgical duration, determined by a quadratic regression fit (R^2^ = 0.6093) of the CUSUM data, was reached after approximately 9 anastomoses **(Figure 4C)**. For surgical skill scores, the CUSUM curve (quadratic fit, R^2^ = 0.8874) indicated proficiency was achieved after approximately 10 anastomoses **(Figure 4D)**. The substantially fewer procedures required to reach proficiency in 5XFAD mice, despite a more moderate R^2^ value for the surgical duration CUSUM fit, underscores the efficacy of the precedent training in SD rats for facilitating skill acquisition in a more challenging supermicrosurgical model.

Of the ten 5XFAD transgenic mice that underwent surgery, one experienced intraoperative mortality attributed to complications with intraperitoneal anesthesia. The remaining nine mice (90% procedural survival) recovered uneventfully and remained in good health at 1 months post-surgery. To assess long-term anastomotic function, a follow-up surgical exploration with ICG lymphangiography was performed in a subset of three surviving mice at two weeks post-surgery, which confirmed sustained patency of the lymph node-venous shunts. These animals were subsequently available for preliminary behavioral assessments, demonstrating the robustness of the LNVA model for future longitudinal studies. The anesthetic-related adverse event has prompted consideration of transitioning to inhalational anesthesia for future procedures to enhance safety.

During ICG lymphangiography via nasal instillation for patency assessment, consideration was given to the potential pathways of tracer distribution, particularly in light of emerging research on intranasal central nervous system drug delivery. A pilot experiment using intranasal Evans Blue dye in a separate cohort revealed staining of the dura mater.^24^ This observation suggests that the visualization of DCLNs following intranasal ICG administration may result from a multi-pathway distribution, **(Figure 2B)** potentially involving dural lymphatic or perivascular clearance routes in addition to direct nasopharyngeal lymphatic uptake, a hypothesis warranting further investigation.

## Discussion

This study successfully establishes and standardizes a lymphatic-venous bypass supermicrosurgical technique, transitioning from an SD rat training model to a clinically relevant 5XFAD AD transgenic mouse application model. Crucially, we have systematically quantified the learning curve of a junior surgeon mastering this complex procedure, integrating UWOMSA scores with CUSUM analysis. Our findings not only validate the efficacy of a staged training paradigm in accelerating skill acquisition and transfer to a more challenging model but also, through exploratory tracer studies, provide preliminary evidence suggesting multi-pathway communication between the nasal cavity, DCLNs, and potentially intracranial structures. These insights offer new perspectives on the intricate connections within the cranio-cervical lymphatic system and highlight potential translational avenues.

The pathogenesis of Alzheimer’s disease (AD) is complex, with impaired clearance of cerebral metabolic waste, such as amyloid-beta (Aβ) protein, recognized as a potential pathological mechanism^6,25–27^. The discovery of the brain’s glymphatic system and meningeal lymphatic vessels has positioned the cervical lymphatic system, particularly the DCLNs, as a key downstream pathway for brain waste and cerebrospinal fluid (CSF) drainage.^28^ Building on this understanding, our group, under the leadership of Prof. Xie, pioneered the concept of surgically enhancing cervical lymphatic drainage via lymphatic-venous bypass to promote cerebral waste clearance and potentially ameliorate AD, initiating groundbreaking clinical explorations.^11^ This innovative strategy has garnered significant international attention within the neuroscience and lymphology communities, with multiple centers actively pursuing related clinical trials.

Despite this considerable clinical potential, a significant translational gap existed due to the extreme technical challenges of performing such demanding supermicrosurgical lymphatic anastomoses in small animals, especially in genetically defined AD transgenic mouse models. Consequently, the field lacked a stable, reproducible “gain-of-function” surgical AD mouse model to simulate this clinical intervention and facilitate mechanistic investigations. The present study successfully bridges this critical preclinical research gap. Through rigorous protocol design, systematic surgeon training, and technical optimization, we have not only established this much-needed model but, more importantly, provided a unique, clinically-relevant preclinical platform for dissecting AD pathophysiology and evaluating novel therapeutic interventions.

Supermicrosurgical techniques(less than 0.8 mm vessels), characterized by their operational intricacy and high degree of difficulty, impose stringent demands on surgeons, typically involving steep learning curves.^29^ This study employed the internationally recognized UWOMSA scoring system, augmented by CUSUM analysis,^30^ to objectively and quantitatively assess the learning trajectory of a junior surgeon mastering lymphatic-venous bypass. The data revealed a classic learning pattern: in the SD rat training phase, surgical duration exhibited a rapid initial decline followed by stabilization, while surgical scores progressively improved. CUSUM analysis indicated that operational proficiency, defined by the stabilization of both time and skill metrics, was achieved after approximately 26-27 anastomoses (equivalent to bilateral procedures in ∼13 SD rats). This quantitative benchmark not only provides an objective standard for evaluating lymphatic-venous bypass skill acquisition but also reaffirms that sufficient, mentored practice is indispensable for cultivating proficiency in complex supermicrosurgical procedures.^27^

The selection of small-bodied (≈50g) SD rats as the initial training model was a deliberate decision with multifaceted advantages. Firstly, while their cervical vascular and lymphatic structures remain significantly larger than those in 5XFAD mice, their relatively smaller operative field and more delicate tissues, compared to standard-weight SD rats, render them an ideal transitional platform from conventional microsurgery to the ultra-fine manipulations required in mice. Secondly, as evidenced by our study, this staged training paradigm yields considerable economic and psychological benefits. Transgenic mouse models are typically expensive and have long breeding cycles. By completing the bulk of the initial learning, technical exploration, and skill consolidation on a cost-effective SD rat model, we significantly reduced early-stage errors and animal attrition in the valuable transgenic cohort, thereby optimizing resource allocation. Concurrently, this progressive approach likely bolstered the surgeon’s confidence when transitioning to the more demanding and less forgiving transgenic mouse surgeries, mitigating performance anxiety and facilitating a more stable application of acquired skills.

The successful transfer of skills to the 5XFAD mouse model further validates the overall efficacy of our two-stage training protocol. In mice, the surgeon achieved the CUSUM curve vertex, indicating proficiency, after only approximately 9-10 anastomoses, demonstrating a markedly accelerated learning and adaptation phase. This clearly indicates that the supermicrosurgical expertise and fine motor skills honed on the rat model were effectively transferable to the anatomically more intricate and technically more demanding mouse model. This finding has significant implications for optimizing training protocols for other advanced surgical techniques, suggesting that comprehensive pre-training on less costly, relatively easier surrogate models can substantially shorten the learning curve and improve success rates when transitioning to more valuable or challenging target models, particularly disease model animals.

Precise morphometric comparison using ImageJ revealed statistically significant differences in EJV diameter and DCLN size between SD rats and 5XFAD mice, with the latter possessing considerably smaller structures. These inherent anatomical disparities directly contribute to the increased operational difficulty of surgery in the mouse model. This necessitates not only superior microsurgical dexterity and finer supermicrosurgical instrumentation but also enhanced hand-eye coordination, stable motor control, and acute three-dimensional spatial perception. For instance, creating a venotomy and achieving precise apposition and suturing of the transected lymph node surface to the EJV wall in mice demands a level of precision far exceeding that required in rats. These challenges underscore why even surgeons with existing microsurgical experience require a distinct adaptation and learning period when first encountering murine surgery. Our findings emphasize the critical importance, when designing similar technology transfer studies or surgical training programs, of fully appreciating the anatomical differences between training and target models and implementing targeted skill reinforcement and mental preparation. In this study, intranasal instillation of indocyanine green (ICG) effectively visualized deep cervical lymph nodes (DCLNs) and confirmed anastomotic patency in both SD rats and 5×FAD mice. Among nine surviving 5×FAD mice, all 18 bilateral anastomoses (100%) demonstrated immediate postoperative patency, with fluorescent signal clearly flowing from the lymph node into the external jugular vein (EJV), supporting the reliability of this model for future longitudinal studies. Traditionally, lymphatic drainage from the nasal mucosa is thought to follow a superficial route, progressing through submandibular and superficial cervical nodes before reaching deep cervical lymph nodes.^32^ However, emerging studies on nasal–brain lymphatic connectivity,^7^ along with our own intraoperative ICG and Evans Blue imaging, suggest an alternative drainage route through the nasopharyngeal lymphatic plexus. In our experiments, intranasal Evans Blue administration resulted in visible staining of the dura near the skull base, while dural injections of Evans Blue also produced retrograde staining patterns near the nasopharynx under mild pressure. These findings imply the existence of a bidirectional or pressure-sensitive nasal–meningeal lymphatic pathway. Importantly, this mucosal or submucosal instillation technique may not only help avoid contamination from local surgical wounds during ICG-based imaging, but also offer a novel strategy for targeted delivery into the CNS. Unlike aerosol-based intranasal delivery systems that rely on passive diffusion or airflow-dependent transport,^33,34^ our method may utilize more direct anatomical conduits—such as perineural pathways or lymphatic networks—for enhanced CNS bioavailability. This could be particularly valuable for delivering macromolecules, gene therapy vectors, or cell-based treatments that struggle to cross the blood–brain barrier. Further studies are warranted to elucidate these transport mechanisms, optimize administration parameters, and evaluate safety and efficacy.

This study has several limitations. The learning curve relied on a single surgeon, limiting generalizability; multi-center studies are needed. Our focus was on establishing the surgical model, with long-term AD pathology assessments being ongoing. Preliminary nasal drainage findings require further investigation with advanced imaging, including in vivo two-photon microscopy^35^ and high-resolution MRI lymphangiography^36^, and quantitative methods. Finally, we acknowledge that achieving complete standardization of suture quality is a key challenge for any procedure that is highly dependent on an operator’s supermicrosurgical skill. To address this, we plan to develop and apply microsurgical robots to this animal model in future work, aiming to overcome technical variability and achieve the highest level of standardization and reproducibility.^37,38^

By systematically deconstructing the supermicrosurgical learning curve, we have created a validated and accessible training paradigm. This key contribution helps to democratize a previously exclusive technique, empowering more research groups to investigate the role of cervical lymphatic drainage in AD pathophysiology. This robust platform, combined with our intriguing preliminary findings on nasal-meningeal pathways, opens new avenues for both mechanistic discovery and the development of novel delivery strategies for AD therapeutics. Ultimately, this work provides a critical translational bridge, accelerating the journey from a promising surgical concept to a potential new class of treatments for Alzheimer’s disease.

## Conclusion

We successfully established a reproducible supermicrosurgical LVA model in a transgenic AD mouse. We demonstrated that a staged training protocol, beginning with a rat model, enables a junior surgeon to achieve proficiency efficiently, as validated by a quantitative learning curve analysis. This accessible and highly patent surgical platform is a critical tool for future preclinical investigation into the role of lymphatic drainage in Alzheimer’s disease.

## Provenance and peer review

Not commissioned, externally peer-reviewed

## Conflicts of Interest

None declared.

## Sources of Funding

This study received no external funding.

## Ethical Approval

Ethical approval was obtained from the Institutional Animal Care and Use Committee. Details withheld for peer review.

## Consent

Not applicable.

## Artificial Intelligence Use

This study followed the TITAN Guidelines 2025 for the responsible use of AI in scientific writing and analysis. Generative AI (ChatGPT, OpenAI GPT-4Oversion) was used solely to enhance language clarity and grammar during manuscript drafting and revision. All AI outputs were critically reviewed and edited by the authors, who take full responsibility for the final content. No scientific interpretation, data analysis, figure generation, or image processing involved the use of AI.

## Notes

### Competing Interest Statement

The authors have declared no competing interest.

## References

1. Iliff JJ, Wang M, Liao Y, et al. A Paravascular Pathway Facilitates CSF Flow Through the Brain Parenchyma and the Clearance of Interstitial Solutes, Including Amyloid β. Sci Transl Med. 2012;4(147). doi:10.1126/scitranslmed.3003748

2. Nedergaard M. Garbage Truck of the Brain. Science. 2013;340(6140):1529–1530. doi:10.1126/science.1240514

3. Louveau A, Smirnov I, Keyes TJ, et al. Structural and functional features of central nervous system lymphatic vessels. Nature. 2015;523(7560):337–341. doi:10.1038/nature14432

4. Nedergaard M, Goldman SA. Glymphatic failure as a final common pathway to dementia. Science. 2020;370(6512):50–56. doi:10.1126/science.abb8739

5. Rasmussen MK, Mestre H, Nedergaard M. The glymphatic pathway in neurological disorders. Lancet Neurol. 2018;17(11):1016–1024. doi:10.1016/S1474-4422(18)30318-1

6. Harrison IF, Ismail O, Machhada A, et al. Impaired glymphatic function and clearance of tau in an Alzheimer’s disease model. Brain. 2020;143(8):2576–2593. doi:10.1093/brain/awaa179

7. Yoon JH, Jin H, Kim HJ, et al. Nasopharyngeal lymphatic plexus is a hub for cerebrospinal fluid drainage. Nature. 2024;625(7996):768–777. doi:10.1038/s41586-023-06899-4

8. Wang L, Zhang Y, Zhao Y, Marshall C, Wu T, Xiao M. Deep cervical lymph node ligation aggravates AD[like pathology of APP/PS1 mice. Brain Pathol. 2019;29(2):176–192. doi:10.1111/bpa.12656

9. Kim K, Abramishvili D, Du S, et al. Meningeal lymphatics-microglia axis regulates synaptic physiology. Cell. Published online March 2025:S0092867425002107. doi:10.1016/j.cell.2025.02.022

10. Du T, Raghunandan A, Mestre H, et al. Restoration of cervical lymphatic vessel function in aging rescues cerebrospinal fluid drainage. Nat Aging. 2024;4(10):1418–1431. doi:10.1038/s43587-024-00691-3

11. Xie Q, Louveau A, Pandey S, Zeng W, Chen WF. Rewiring the Brain – the Next Frontier in Supermicrosurgery. Plast Reconstr Surg. Published online July 18, 2023. doi:10.1097/PRS.0000000000010933

12. Hong JP, Chen WF, Nguyen DH, Xie Q. A Proposed Role for Lymphatic Supermicrosurgery in the Management of Alzheimer’s Disease: A Primer for Reconstructive Microsurgeons. Arch Plast Surg. Published online January 10, 2025:a-2513-4313. doi:10.1055/a-2513-4313

13. Lu H, Tan Y, Xie Q. Preliminary observation of deep cervical lymphatic-venous anastomosis under off-eyepiece 3D microscope to treat an elderly patient with cognitive impairment. Chin J Microsurg. Published online 2022:570–574.

14. Li X, Zhang C, Fang Y, et al. Promising outcomes 5 weeks after a surgical cervical shunting procedure to unclog cerebral lymphatic systems in a patient with Alzheimer’s disease. Gen Psychiatry. 2024;37(3):e101641. doi:10.1136/gpsych-2024-101641

15. Chen JY, Zhao DW, Yin Y, et al. Deep cervical lymphovenous anastomosis (LVA) for Alzheimer’s disease microsurgical procedure in a prospective cohort study. Int J Surg. Published online May 20, 2025. doi:10.1097/js9.0000000000002490

16. Xu G, Zhang Q, Li Y, et al. Cerebrospinal Fluid p-Tau181/Amyloid Beta 42 Ratio Identifies Lymphatic-Venous Anastomosis Patients Who Respond to and Benefit from the Surgery for Relief of Cognitive Impairment with a Diagnostic Accuracy of 0.744. ACS Omega. Published online July 16, 2025. doi:10.1021/acsomega.5c03650

17. Fang R, Jin L, Lu H, Xie Q, Yang X, Kueckelhaus M. A Novel Microsurgical Model of Cervical Lymph Node-to-Vein Anastomosis (LNVA) for Studying Brain Lymphatic Outflow. J Craniofac Surg. Published online June 18, 2025. doi:10.1097/scs.0000000000011535

18. Oakley H, Cole SL, Logan S, et al. Intraneuronal β-Amyloid Aggregates, Neurodegeneration, and Neuron Loss in Transgenic Mice with Five Familial Alzheimer’s Disease Mutations: Potential Factors in Amyloid Plaque Formation. J Neurosci. 2006;26(40):10129–10140. doi:10.1523/JNEUROSCI.1202-06.2006

19. Improving bioscience research reporting: The ARRIVE guidelines for reporting animal research -Carol Kilkenny, William J. Browne, Innes C. Cuthill, Michael Emerson, Douglas G. Altman, 2010. Accessed May 31, 2025. https://journals.sagepub.com/doi/abs/10.4103/0976-500X.72351

20. Temple CLF, Ross DC. A New, Validated Instrument to Evaluate Competency in Microsurgery: The University of Western Ontario Microsurgical Skills Acquisition/Assessment Instrument [Outcomes Article]: Plast Reconstr Surg. 2011;127(1):215–222. doi:10.1097/PRS.0b013e3181f95adb

21. Proulx ST, Luciani P, Derzsi S, et al. Quantitative Imaging of Lymphatic Function with Liposomal Indocyanine Green. Cancer Res. 2010;70(18):7053–7062. doi:10.1158/0008-5472.CAN-10-0271

22. Pak CS, Suh HP, Kwon JG, Cho MJ, Hong JP. Lymph Node to Vein Anastomosis (LNVA) for lower extremity lymphedema. J Plast Reconstr Aesthet Surg. 2021;74(9):2059–2067. doi:10.1016/j.bjps.2021.01.005

23. Bailey EA, Pandey SK, Chen WF. Advances in Surgical Lymphedema Management: The Emergence and Refinement of Lymph Node-to-Vein Anastomosis (LNVA). Curr Surg Rep. 2024;12(5):83–88. doi:10.1007/s40137-024-00395-y

24. Maloveska M, Danko J, Petrovova E, et al. Dynamics of Evans blue clearance from cerebrospinal fluid into meningeal lymphatic vessels and deep cervical lymph nodes. Neurol Res. 2018;40(5):372–380. doi:10.1080/01616412.2018.1446282

25. Mentis AFA, Dardiotis E, Chrousos GP. Apolipoprotein E4 and meningeal lymphatics in Alzheimer disease: a conceptual framework. Mol Psychiatry. 2021;26(4):1075–1097. doi:10.1038/s41380-020-0731-7

26. Boland B, Yu WH, Corti O, et al. Promoting the clearance of neurotoxic proteins in neurodegenerative disorders of ageing. Nat Rev Drug Discov. 2018;17(9):660–688. doi:10.1038/nrd.2018.109

27. Ye C, Wang S, Niu L, et al. Unlocking potential of oxytocin: improving intracranial lymphatic drainage for Alzheimer’s disease treatment. Theranostics. 2024;14(11):4331–4351. doi:10.7150/thno.98587

28. Ma Q, Ineichen BV, Detmar M, Proulx ST. Outflow of cerebrospinal fluid is predominantly through lymphatic vessels and is reduced in aged mice. Nat Commun. 2017;8(1):1434. doi:10.1038/s41467-017-01484-6

29. Hong JP (Jp), Song S, Suh HSP. Supermicrosurgery: Principles and applications. J Surg Oncol. 2018;118(5):832–839. doi:10.1002/jso.25243

30. Yap C, Colson ME, Watters DA. CUMULATIVE SUM TECHNIQUES FOR SURGEONS: A BRIEF REVIEW. ANZ J Surg. 2007;77(7):583–586. doi:10.1111/j.1445-2197.2007.04155.x

31. Hong JP, Masoodi Z, Tzou CHJ. Attributes of a Good Microsurgeon—A Brief Counsel to the Up-and-Coming Prospects. Arch Plast Surg. 2023;50(01):130–140. doi:10.1055/s-0042-1759786

32. Fernández JMS, Santaolalla F, Rey ASD, Martínez-Ibargüen A, González A, Iriarte MR. Preliminary study of the lymphatic drainage system of the nose and paranasal sinuses and its role in detection of sentinel metastatic nodes. Acta Otolaryngol (Stockh). 2005;125(5):566–570. doi:10.1080/00016480510036457

33. Fonseca LC, Lopes JA, Vieira J, et al. Intranasal drug delivery for treatment of Alzheimer’s disease. Drug Deliv Transl Res. 2021;11(2):411–425. doi:10.1007/s13346-021-00940-7

34. Meredith ME, Salameh TS, Banks WA. Intranasal Delivery of Proteins and Peptides in the Treatment of Neurodegenerative Diseases. AAPS J. 2015;17(4):780–787. doi:10.1208/s12248-015-9719-7

35. Miller MJ, Wei SH, Parker I, Cahalan MD. Two-Photon Imaging of Lymphocyte Motility and Antigen Response in Intact Lymph Node. Science. 2002;296(5574):1869–1873. doi:10.1126/science.1070051

36. Forte AJ, Boczar D, Huayllani MT, et al. Use of magnetic resonance imaging lymphangiography for preoperative planning in lymphedema surgery: A systematic review. Microsurgery. 2021;41(4):384–390. doi:10.1002/micr.30731

37. Wessel KJ, Dahmann S, Kueckelhaus M. Expanding Applications and Future of Robotic Microsurgery. J Craniofac Surg. 2025;36(1):367. doi:10.1097/SCS.0000000000010860

38. Kueckelhaus M, Nistor A, van Mulken T, et al. Clinical experience in open robotic-assisted microsurgery: user consensus of the European Federation of Societies for Microsurgery. J Robot Surg. 2025;19(1):171. doi:10.1007/s11701-025-02338-w

